# Adipocyte metabolic state regulates glial phagocytic function

**DOI:** 10.1101/2024.09.24.614765

**Authors:** Mroj Alassaf, Akhila Rajan

**Affiliations:** Basic Sciences Division, Fred Hutch, Seattle, WA-98109. The USA

## Abstract

**Highlights:**

- Prolonged exposure to an obesogenic diet result in a starvation-like metabolic response in adipose tissue.
- Obesogenic diet-induced mitochondrial lipid catabolism in adipose tissue impacts glial phagocytic function.
- Adipocyte ApoB is a novel regulator of glial phagocytic function.
- LpR1, on ensheathing glia, is required for glial response to axonal injury.

Obesity and type 2 diabetes are well-established risk factors for neurodegenerative disorders^1–4^, yet the underlying mechanisms remain poorly understood. The adipocyte-brain axis is crucial for brain function, as adipocytes secrete signaling molecules, including lipids and adipokines, that impinge on neural circuits to regulate feeding and energy expenditure^5^. Disruptions in the adipocyte-brain axis are associated with neurodegenerative conditions^6^, but the causal links are not fully understood. Neural debris accumulates with age and injury, and glial phagocytic function is crucial for clearing this debris and maintaining a healthy brain microenvironment^7–9^. Using adult *Drosophila,* we investigate how adipocyte metabolism influences glial phagocytic activity in the brain. We demonstrate that a prolonged obesogenic diet increases adipocyte fatty acid oxidation and ketogenesis. Genetic manipulations that mimic obesogenic diet-induced changes in adipocyte lipid and mitochondrial metabolism unexpectedly reduce the expression of the phagocytic receptor Draper in *Drosophila* microglia-like cells in the brain. We identify *Apolpp*—the *Drosophila* equivalent of human apolipoprotein B (ApoB)—as a critical adipocyte-derived signal that regulates glial phagocytosis. Additionally, we show that Lipoprotein Receptor 1 (LpR1), the LDL receptor on phagocytic glia, is required for glial capacity to clear injury-induced neuronal debris. Our findings establish that adipocyte-brain lipoprotein signaling regulates glial phagocytic function, revealing a novel pathway that links adipocyte metabolic disorders with neurodegeneration.

Graphical abstract

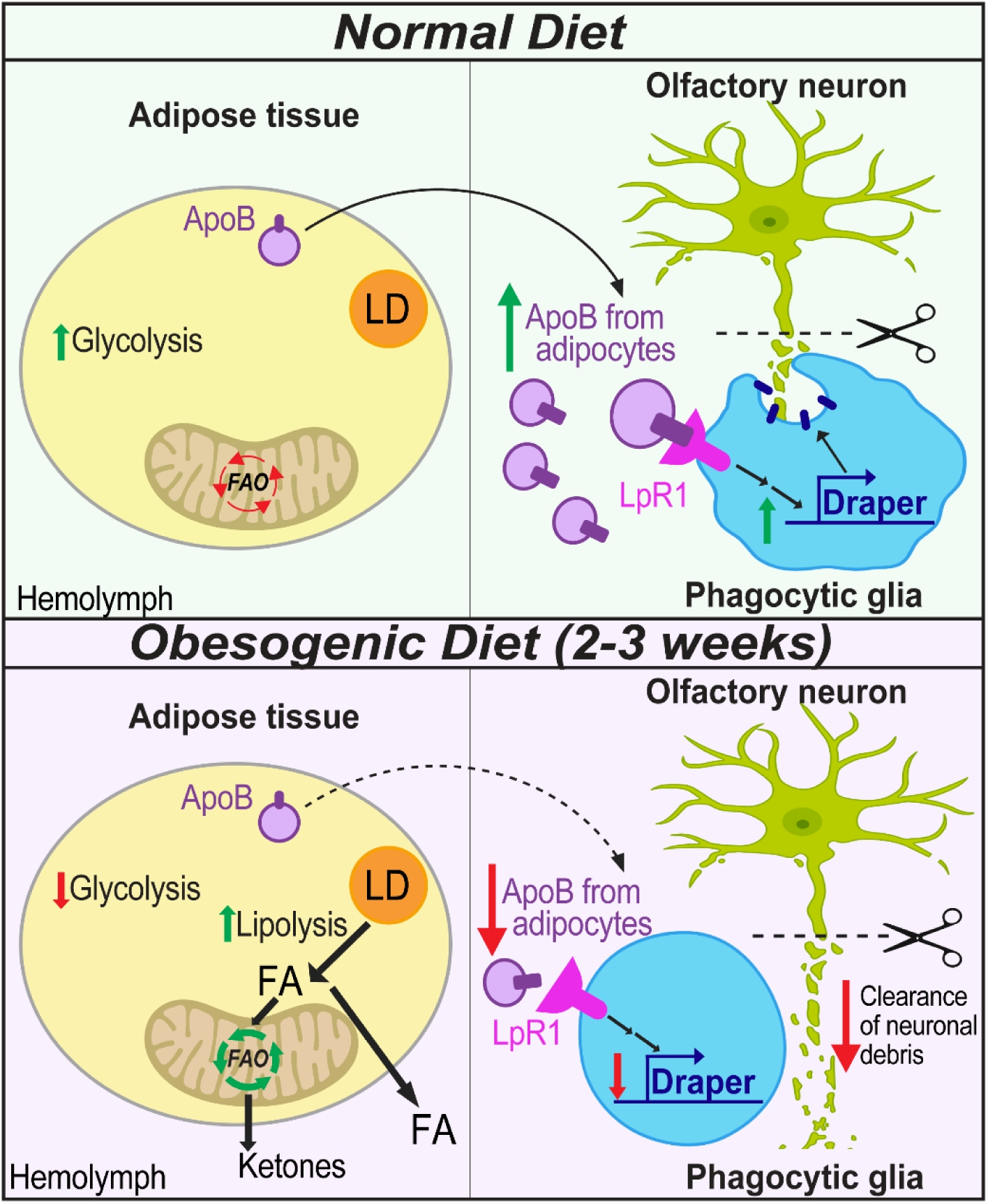

## Introduction

Obesity and type 2 diabetes are increasingly recognized as significant risk factors for the development of dementia and neurodegenerative disorders ^1–4^. While the mechanistic link between these conditions is not fully elucidated, a common feature is the disruption of the adipose tissue-brain axis ^10–13^. Once regarded primarily as a passive energy depot, adipose tissue has emerged as a multifaceted endocrine organ that actively secretes diverse signaling molecules, profoundly influencing brain function ^5^. These adipocyte-derived signals can impact various processes such as neuroinflammation, oxidative stress, and synaptic plasticity^11,13,14^. Therefore, understanding the mechanisms governing adipose tissue-brain cross-talk holds promise for identifying novel therapeutic strategies to combat neurodegeneration.

The brain, rich in lipids yet poor in lipid synthesis^15^, relies on peripheral tissues to supply necessary lipids to fuel its massive energy demands ^16,17^. Adipose tissue, as the primary lipid reservoir, plays a crucial role in this supply chain, and disruptions in its lipid homeostasis can adversely affect brain function ^18^. Lipoprotein particles are specialized vesicles consisting of a phospholipid monolayer surrounding a core of esterified cholesterol and triglycerides (TAG) that facilitate lipid trafficking between cells and organs. Apolipoproteins scaffold lipoproteins and act as chaperones to deliver lipids to specific locations. Apolipoprotein B (ApoB) is the primary apolipoprotein responsible for transporting lipids from the periphery into the brain. ApoB can traverse the restrictive blood-brain barrier by binding to its low-density lipoprotein receptor (LDLR) receptor ^19–22^. While there is abundant evidence of dysfunctional ApoB signaling in obesity, type 2 diabetes ^23–25^, and neurodegenerative disorders ^26,27^, a direct causal link involving ApoB has yet to be investigated.

*Drosophila* exhibits remarkable similarities to mammals in its lipid metabolism. They express the ApoB ortholog Apolpp, produced in adipocytes and chaperones lipophorins (*ApoB-Lpp*) that carry most lipids in circulation^28–31^. ApoB-Lpp provides lipids to developing organs during *Drosophila* development^32^ and delivers lipids to the *Drosophila* brain^33^. Depletion of ApoB in adipocytes significantly reduces brain lipid stores, particularly TAG^33^. Furthermore, *Drosophila* lipoprotein receptors, LpR1 and LpR2, resemble mammalian LDLRs^34,35^ and facilitate Lpp internalization in peripheral tissue^32,36^. Also, similar to the mammalian brain, neuron-glia lipid shuttling within the *Drosophila* brain involves GLaZ/ApoD with Glaz/ApoD overexpression, providing neuroprotection by enabling glia to accumulate neutral lipids and protect against ROS-induced cell death^37^. Recent studies indicate that ApoB-Lpp delivery to the brain influences systemic insulin signaling and feeding behavior in *Drosophila*^38–40^. However, the impact of ApoB-Lpps on glial function and possible neuroprotection remains unclear.

A hallmark of neurodegenerative disorders is the accumulation of protein aggregates and cellular debris, creating a toxic cellular environment and causing secondary neuronal death ^41^. Microglia, the brain’s resident macrophages, mitigate this damage by phagocytosing the debris, thereby maintaining a healthy cellular environment ^42^. In *Drosophila*, ensheathing glia performs functions like those of microglia^8^. They detect neuronal debris using the engulfment receptor Draper, which is homologous to mammalian multiple epidermal growth factor-like domain protein 10 (MEGF10)^43,44^. Upon recognizing debris, ensheathing glia extend their membranes to encase and phagocytose the debris ^8,45,46^. Several studies have demonstrated that Draper signaling in ensheathing glia, like MEGF10 signaling in microglia, mediates the clearance of apoptotic cells and cellular debris in the *Drosophila* nervous system ^47–50^. Notably, studies using *Drosophila* models of Alzheimer’s disease have demonstrated that ensheathing glia efficiently clear transgenically expressed human amyloid beta (Aβ) through Draper-mediated phagocytosis^51^, mirroring the function of microglia in mammalian models of Alzheimer’s^52^. Furthermore, aged flies exhibit declining glial phagocytic function like that observed in the aging human brain ^48^. These findings highlight the conservation of this mechanism across species.

We have previously shown that an obesogenic diet (high sugar diet; HSD) disrupts lipid homeostasis and causes adipocyte insulin resistance ^39^. Furthermore, we have shown that HSD induces glial insulin resistance, leading to impaired clearance of neuronal debris by disrupting Draper signaling ^53^. Given the conservation of adipocyte-brain signals^54,55^, metabolic processes^56^, and glial biology^45,57^ between flies and mammals, we investigate how the metabolic state of adipocytes on normal versus obesogenic diets influences glial health and explore the potential role of adipocyte ApoB lipophorins in modulating glial function, with implications for neurodegenerative disorders.

Here, we unravel a novel adipocyte-glial metabolic coupling that may underlie the obesity-associated heightened risk of neurodegenerative disorders. We directly link adipocyte mitochondrial metabolism and Draper-mediated glial phagocytic function. We show that HSD-induced changes in adipocyte lipolysis, mitochondrial morphology, metabolism, and redox state remotely regulate glial Draper levels. We identify two specific mechanisms by which HSD-induced adipocyte dysfunction impacts glial biology. First, we show that HSD-induced mitochondrial changes and increased ketogenesis disrupt glial Draper signaling. Second, we show that intact adipocyte-brain lipoprotein signaling is required for ensheathing glia’s response to axonal injury. These findings illuminate a critical link between adipocyte dysfunction and glial behavior, suggesting potential pathways by which obesity may increase the risk of neurodegenerative disorders and highlighting novel therapeutic targets for intervention.

## Results

### Prolonged exposure to high-sugar diets (HSD) shifts adipocyte metabolism towards mitochondrial oxidative phosphorylation

Cellular energy production relies on balancing two major metabolic pathways: Glycolysis and Oxidative Phosphorylation (OxPhos)^58^. We examined the effects of a high-sugar diet (HSD) on glycolysis in adult *Drosophila* adipocytes. Pyruvate kinase (*pyk*) catalyzes the final step of glycolysis, converting phosphoenolpyruvate (PEP) to pyruvate while generating ATP (Figure 1A). Since converting PEP to pyruvate is an irreversible process, it determines glycolysis flux, making it a key control point in the pathway^59,60^. Loss of *pyk* severely impairs glycolysis in *Drosophila* and results in the accumulation of glycolytic intermediates^61^. Given that *pyk* is a rate-limiting enzyme, we monitored *pyk* expression levels by isolating RNA from abdominal segments of adult flies fed either a normal diet (ND) or HSD for three weeks and performing qPCR for *pyk*. We observed that 3-weeks of HSD treatment significantly downregulated *pyk* expression, indicating reduced glycolysis (Figure 1B).

**Figure 1.**
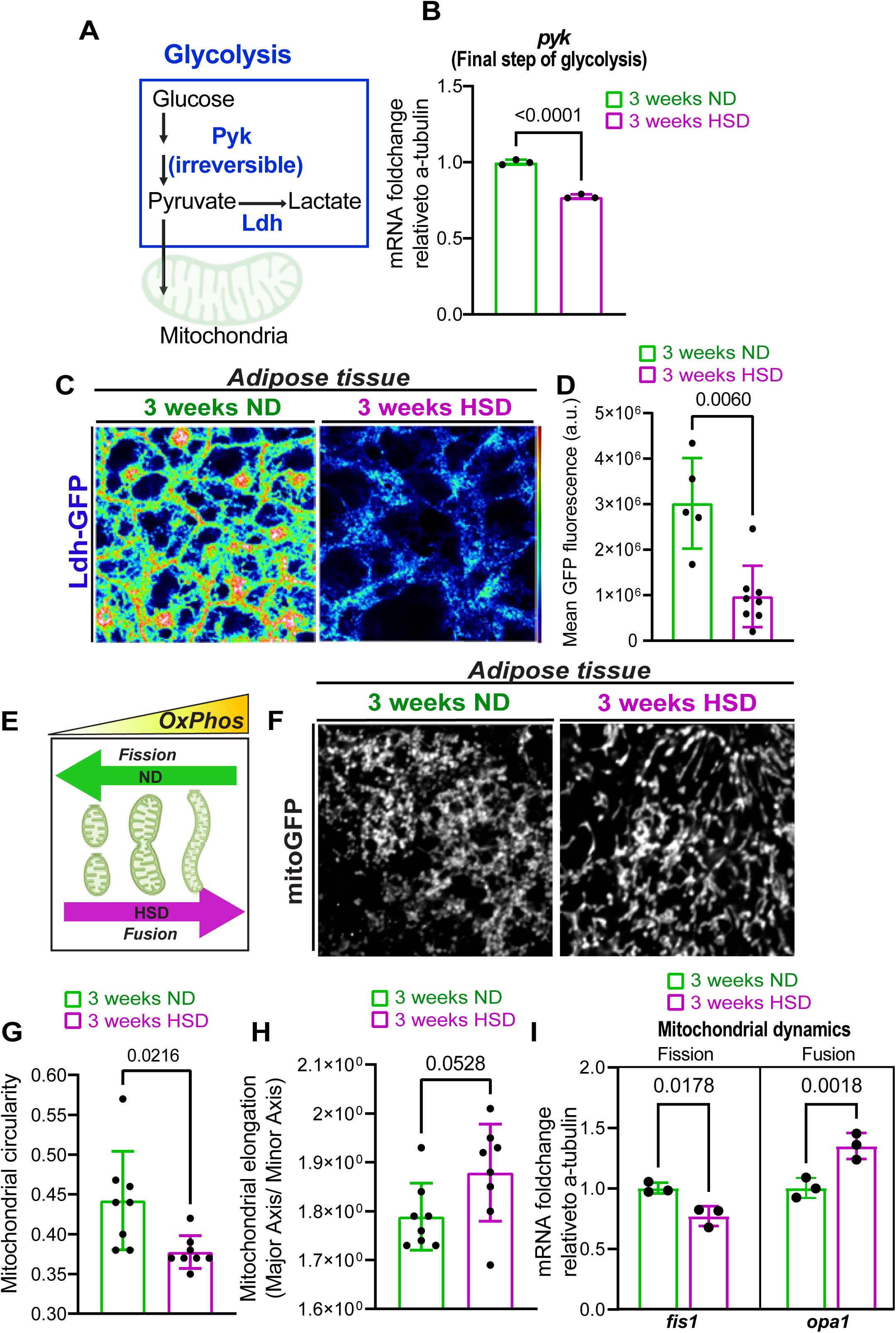
A high-sugar diet (HSD) induces a metabolic shift in adipocytes from glycolysis to oxidative phosphorylation (OxPhos). A) Schematic summarizing the cytosolic glycolysis pathway. Pyruvate kinase (pyk) irreversibly catalyzes the final step of glycolysis, converting phosphoenolpyruvate (PEP) to pyruvate. Ldh catalyzes the conversion of pyruvate to lactate. B) Mean fold change in pyk mRNA levels relative to alpha-tubulin in the adipose tissue of flies fed either an ND or an HSD for 3 weeks. Student’s t-test with Welch’s correction. N = 3 technical replicates of cDNA collected from 30 fly abdominal segments/treatment. C) Confocal images of Ldh-GFP in the adipose tissue of flies fed either an ND or an HSD for 3 weeks. Scale bar=20um. D) Mean Ldh-GFP fluorescent intensity, obtained from Z-stack summation projections of adipose tissue in ND and HSD-fed flies. Student’s t-test with Welch’s correction. N = each circle represents an individual fly. E) Schematic showing the relationship between mitochondrial morphology and oxidative capacity. Elongated mitochondria have higher OxPhos compared to circular mitochondria. F) Confocal images of GFP-tagged mitochondria (mito-GFP) in the fatty tissue of ND and HSD-fed flies. Scale bar=20um. G) Mean mitochondrial circularity and (H) elongation measured from Z-stack max intensity projections of mito-GFP in the adipose tissue of flies fed either an ND or an HSD for 3 weeks. Student’s t-test. N each circle represents an individual fly. I) Mean fold change in the mRNA levels of fis1 and opa1 relative to alpha-tubulin in the adipose tissue of flies fed either an ND or an HSD for 3 weeks. Student’s t-test with Welch’s correction. N = 3 technical replicates of cDNA collected from 30 fly abdominal segments/treatment.

To corroborate this finding, we used a transgenic reporter line expressing GFP-tagged Lactate dehydrogenase (Ldh), another glycolytic enzyme^53,62,63^. Ldh catalyzes the conversion of pyruvate to lactate (Figure 1A), indicating glycolytic activity^64^. Immunohistochemistry revealed that three weeks of HSD treatment reduced Ldh levels in adipose tissue (Figure 1C-D), consistent with decreased *pyk* mRNA levels. Notably, Ldh levels declined in adipose tissue after two weeks of HSD (Figure S1). In contrast, we have previously reported that analysis of the whole brain showed that glycolysis was not significantly depressed at the 2-week exposure timepoint^53^. Overall, these results suggested that HSD exposure reduces adipose tissue glycolysis before such an effect can be observed in the brain^53^.

When glycolysis is impaired, mitochondria rapidly expand to compensate for decreased energy levels^58,65,66^. Consequently, mitochondrial morphology can indicate their activity, with elongated mitochondria having higher OxPhos capacity than circular ones^67,68^ (Figure 1E). We investigated whether the HSD-induced reduction in glycolysis resulted in elongated mitochondria in HSD-fed flies. To this end, we expressed a mitochondrion-targeted GFP (mito-GFP) in adipose tissue using an adipocyte-specific promoter (Lpp) and analyzed mitochondrial morphology. We observed a clear shift towards more elongated and less circular mitochondria (Figure 1F-H). Consistent with this, HSD-fed flies showed downregulation of the mitochondrial fission gene, mitochondrial fission 1 (fis1), and upregulation of the mitochondrial fusion gene, mitochondrial dynamin-like GTPase (opa1) (Figure 1I). Our findings suggest that prolonged (2-3 week) HSD exposure shifts adipose tissue metabolism from glycolysis towards mitochondrial activity, accompanied by changes in mitochondrial morphology to support this metabolic shift.

### HSD Upregulates Adipocyte Mitochondrial Fatty Acid Oxidation

Mitochondria are central to regulating adipocyte metabolism^69^. During nutritional abundance, cells primarily use glycolysis to metabolize glucose, facilitated by insulin signaling, which promotes glucose uptake and storage as triglycerides^70^. In contrast, under metabolic stress with limited glucose availability, mitochondria adapt by expanding and shifting to fatty acid oxidation (FAO)^71^. In obesity and type 2 diabetes, prolonged high circulating insulin levels induce insulin resistance in adipocytes, disrupting lipid homeostasis and increasing lipolysis and FAO^72,73^. Glycolysis and fatty acid oxidation (FAO) are two independent metabolic processes that generate Acetyl-CoA, the primary fuel for the mitochondrial TCA cycle, essential for OxPhos^74^ Figure 2A) under low insulin signaling, such as insulin resistance, cellular glucose uptake decreases, reducing glycolysis and increasing FAO to compensate. Given that HSD decreases glycolysis (Figure 1B-D) and insulin sensitivity^53^ in adipose tissue, we hypothesized a compensatory increase in FAO to maintain Acetyl-CoA levels.

**Figure 2.**
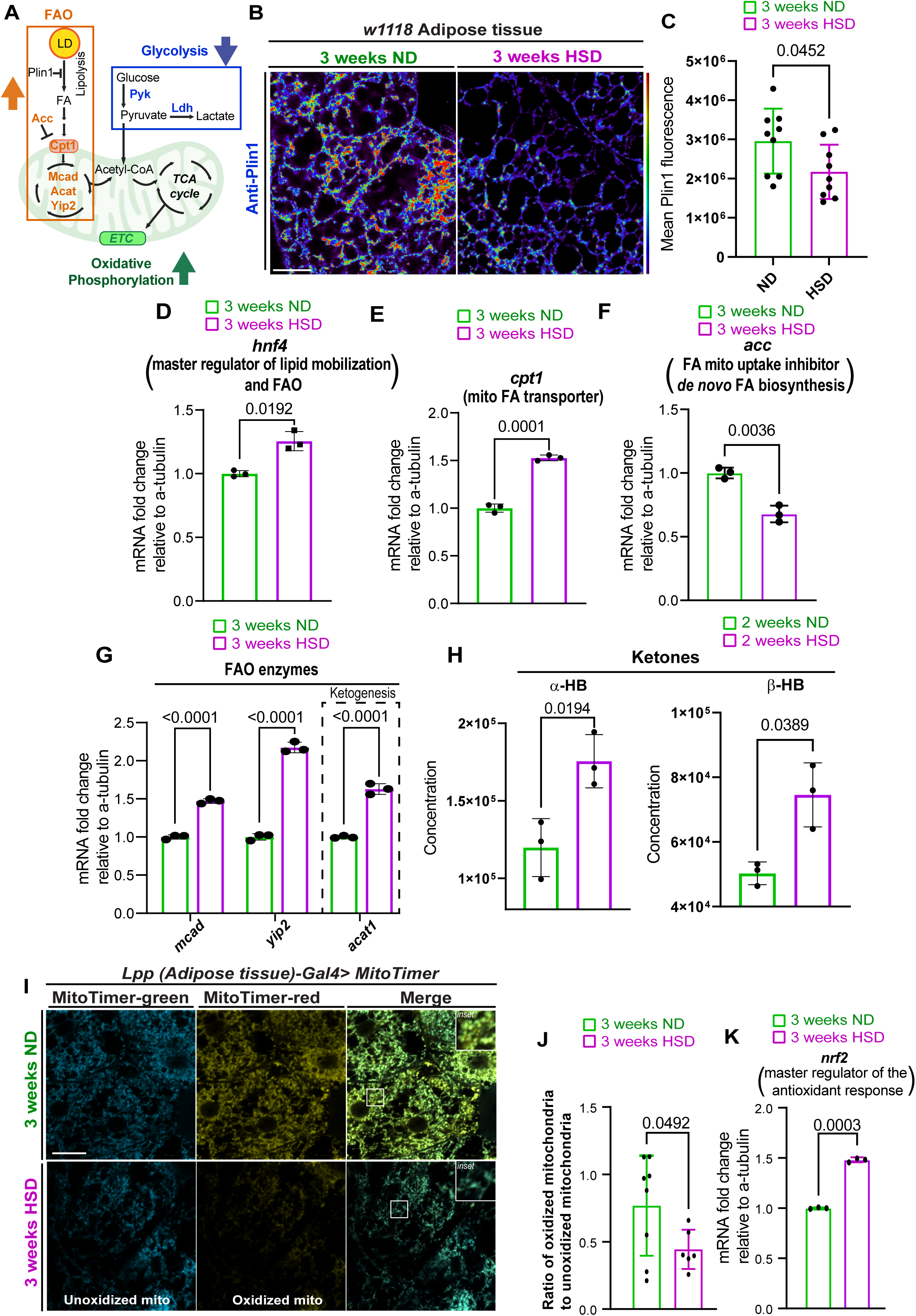
HSD upregulates lipid mobilization and FAO. A) Schematic summarizing the metabolic changes in adipocytes caused by HSD. HAS attenuates glycolysis, leading to a compensatory surge in FAO. B) Confocal images of adipose tissue from w1118 flies fed either an ND or an HSD for 3 weeks, immunostained for Plin1. Student’s t-test. N each circle represents an individual fly. C) Mean Plin1 fluorescent intensity, obtained from Z-stack summation projections of adipose tissue in ND and HSD-fed flies. Student’s T-test with Welch’s correction. N = each circle represents an individual fly. D-G) Mean fold change in the mRNA levels of (D) hnf4, (E) cpt1, (F) acc, and (G) several FAO enzymes relative to alpha-tubulin in the adipose tissue of flies fed either a ND or a HSD for 3 weeks. Student’s t-test with Welch’s correction (D-F) and One-Way ANOVA (G). N = 3 technical replicates of cDNA collected from 30 fly abdominal segments/treatment. H) Mean concentration of the ketone bodies alpha-hydroxybutyrate and beta-hydroxybutyrate in whole flies fed either an ND or a HSD for 2 weeks. Student’s t-test with Welch’s correction. N = 3 technical replicates of 10 flies/replicate. I) Confocal images of adipose tissue from flies expressing MitoTimer after 3 weeks of either ND or HSD. Green fluorescence represents unoxidized mitochondria, while red fluorescence represents oxidized mitochondria. J) Mean ratio of oxidized mitochondria (red channel) to unoxidized mitochondria (green channel) in the adipose tissue of ND and HSD-fed flies. Student’s t-test with Welch’s correction. N = each circle represents an individual fly. K) Mean fold change in nrf2 mRNA levels relative to alpha-tubulin in the adipose tissue of flies fed either an ND or an HSD for 3 weeks. Student’s t-test with Welch’s correction. N = 3 technical replicates of cDNA collected from 30 fly abdominal segments/treatment.

Perilipin 1 (Plin1), known as Lsd1 in flies^75,76^, is a lipid droplet-associated protein that inhibits lipolysis and regulates the release of fatty acids essential for FAO^77,78^. We assessed Plin1 levels in the adipose tissue of ND and HSD-fed flies using immunohistochemistry^79^. HSD-fed flies exhibited reduced Plin1 levels, indicating decreased Plin1-mediated inhibition of lipolysis (Figure 2B-C). Additionally, we used qPCR to analyze the expression of FAO-related enzymes in adipose tissue. HSD upregulated the transcriptional master regulator of lipid mobilization and FAO, hepatocyte nuclear factor 4 (*hnf4*) (Figure 2D), and carnitine palmitoyltransferase 1 (*cpt1*), a key enzyme facilitating fatty acid transport into mitochondria. The fly ortholog is *withered* (*whd*)^80^ - referred to henceforth as Cpt1) (Figure 2E). Conversely, HSD downregulated Acetyl-CoA carboxylase (*acc*), which controls malonyl-CoA, an inhibitor of Cpt1, thus modulating mitochondrial fatty acid uptake (Figure 2F). Furthermore, several mitochondrial FAO enzymes, including acetoacetyl-CoA thiolase (*acat1*), were upregulated in HSD-fed flies (Figure 2G). Acat1 catalyzes the first step of ketogenesis. In support of elevated ketogenesis in the HSD-fed flies, metabolomic analysis revealed increased alpha-hydroxybutyrate (α-HB) and beta-hydroxybutyrate (β-HB), the most abundant ketone bodies in whole flies after 2 weeks of HSD (Figure 2H) and in circulation after one week of HSD (Figure S2) Together, these findings suggest that HSD shifts the adipose tissue metabolic profile to favor FAO.

We hypothesized the HSD-induced mitochondrial expansion and increase in OxPhos (Figure 1) driven by FAO (Figure 2A-H) could elevate the production of reactive oxygen species (ROS), a byproduct of OxPhos. When produced in excess, ROS can cause oxidative stress. To test our hypothesis, we used flies that express MitoTimer, which has been used to evaluate mitochondrial ROS by other independent studies^81,82^. MitoTimer encodes a mitochondria-targeted green fluorescent protein that shifts irreversibly to red fluorescence when oxidized. We subjected the Mito timer reporter flies to an HSD for 3 weeks. Upon evaluating the ratio of oxidized to unoxidized mitochondria, we found a lower red (oxidized) to green (unoxidized) mitochondria ratio in HSD-fed flies, indicating reduced ROS production (Figure 2I-J). We reasoned that this reduction may result from an adaptive antioxidant response. In support of this, β-HB, which is elevated in the HSD flies (Figure 2H & Figure S2), is known to increase antioxidant levels^83^. Indeed, qPCR analysis of adipose tissue showed elevated levels of nuclear factor erythroid 2-related factor 2 (*nrf2*), a master regulator of the antioxidant response, in HSD-fed flies (Figure 2K). These results suggest that HSD-induced mitochondrial expansion and increased FAO reduce ROS levels in adipose tissue, likely by inducing an adaptive antioxidant response.

### Adipocyte mitochondrial and lipid metabolism remotely impacts the glial phagocytic state

We have previously shown that HSD impairs ensheathing glial phagocytic function by inducing glial insulin resistance, downregulating the engulfment receptor Draper^53^. In this study, we observed that HSD alters mitochondrial metabolism and disrupts lipid homeostasis in adipose tissue (Figure 1-2). Given that glia utilizes lipids for energy metabolism^84^, we reasoned that adipocyte-glia signaling likely impacts glial phagocytic function. To test this, we genetically manipulated several key genes involved in mitochondrial and lipid metabolism in adipocytes. We assessed the basal glial phagocytic state by measuring the expression of Draper as a readout.

As reported in the prior section, prolonged obesogenic diets increase ketogenesis due to increased fatty acid (FA) availability (Figure. 2G, H). Hence, we sought to genetically manipulate FA availability and test the effect on glial phagocytic receptor levels. Plin1 is a protein that restricts lipolysis, reducing FA levels. On prolonged HSD, we observed that Plin1 expression reduced, correlating with increased FA levels (Figure 2B-C). Hence, we asked whether the converse state, adipocyte-specific upregulation of Plin1, should reduce FA availability and impact the glial phagocytic receptor expression. To this end, we overexpressed a human Plin1 transgene in flies^76^ and assessed basal draper expression in the ensheathing glia compared to a control transgene. We observed that increasing Plin1 expression in adipocytes correlates with increased glial Draper expression (Figure 3B-D). Overall, this result is consistent with the interpretation that adipocyte FA availability negatively regulates draper expression. Next, we assessed whether increasing intracellular FA levels in adipocytes will have the opposite impact on glial draper expression. In mice, liver-specific knockdown of *CPT1A* interferes with mitochondrial FA uptake and increases hepatic lipid accumulation^85^. Therefore, we performed adipocyte-specific RNAi-mediated knockdown of *Cpt1* in *Drosophila* to assess the impact of increased adipocyte cytosolic FA accumulation on glial draper levels. We observed that flies with adipocyte-specific *Cpt1* knockdown displayed lower draper expression in the ensheathing glia (Figure 3E-G). The genetic manipulations of FA accumulation in adipocytes suggest that FA availability within the adipocyte is inversely correlated with the basal glial phagocytic state.

**Figure 3.**
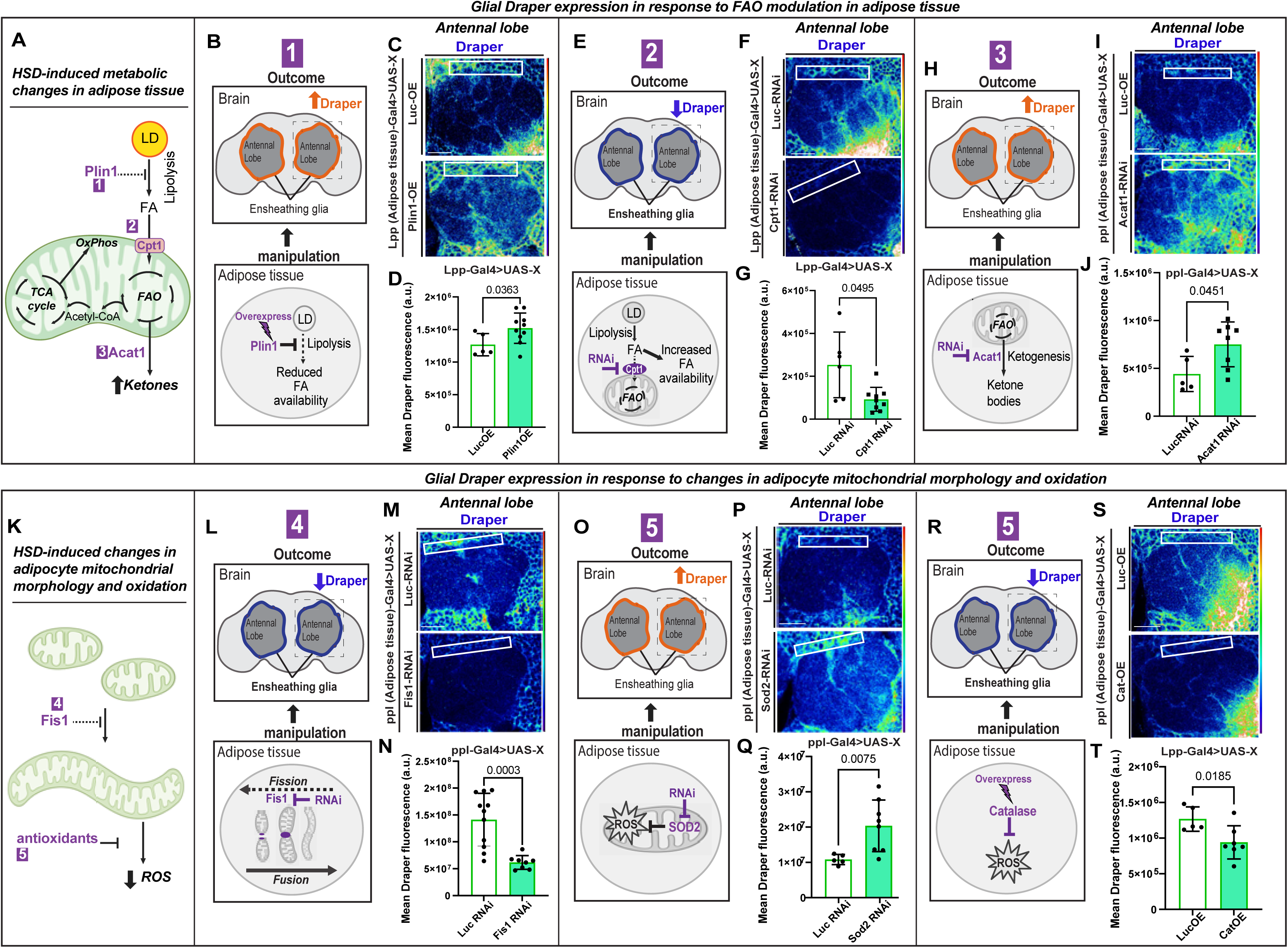
Adipocyte mitochondrial metabolism and morphology impact glial Draper expression. A) Schematic summarizing the pathway of mitochondrial fatty acid oxidation. HSD increases lipolysis by reducing Plin1 inhibition, enhances mitochondrial fatty acid uptake through upregulation of Cpt1, and boosts ketone production via upregulated Acat1. The numbers indicate sites of genetic manipulation performed in B-J. B) Schematic for experimental design and summarized results of C and D. Overexpression of Plin1 in adipose tissue reduces FA availability and increases Draper expression in ensheathing glia. The dotted box highlights the brain region displayed in C. C) Confocal images of the antennal lobe region of flies overexpressing either Luciferase or Plin1 specifically in their adipose tissue immunostained with anti-Draper. D) The Mean fluorescent intensity of Draper measured within a region of interest (white box) coincides with the ensheathing glia’s location. These measurements were derived from a Z-stack summation projection covering the entire depth of the antennal lobe. Student t-test with Welch’s correction. N = each circle represents an individual fly. E) Schematic for experimental design and summarized results of F and G. Knocking down Cpt1 in adipose tissue increases FA availability and reduces Draper expression in ensheathing glia. The dotted box highlights the brain region displayed in C. F) Confocal images of the antennal lobe region of flies expressing either Luciferase-RNAi or Cpt1-RNAi specifically in their adipose tissue immunostained with anti-Draper. G) Mean fluorescent intensity of Draper measured within a region of interest (white box) that coincides with the location of ensheathing glia. These measurements were derived from a Z-stack summation projection covering the entire depth of the antennal lobe. Student t-test with Welch’s correction. N = each circle represents an individual fly. H) Schematic for experimental design and summarized results of I and J. Knocking down the ketogenesis enzyme Acat1 in adipose tissue increases Draper expression in ensheathing glia. The dotted box highlights the brain region displayed in C. I) Confocal images of the antennal lobe region of flies expressing either Luciferase-RNAi or Acat1-RNAi specifically in their adipose tissue immunostained with anti-Draper. J) Mean fluorescent intensity of Draper measured within a region of interest (white box) that coincides with the location of ensheathing glia. These measurements were derived from a Z-stack summation projection covering the entire depth of the antennal lobe. Student t-test with Welch’s correction. N = each circle represents an individual fly. K) Schematic summarizing the effects of HSD on mitochondrial morphology and redox state. HSD causes mitochondrial elongation via the downregulation of Fis1 and reduced ROS by upregulating antioxidants. The numbers indicate sites of genetic manipulation performed in L-T. L) Schematic for experimental design and summarized results of M and N. Knocking down Fis1 in adipose tissue leads to a reduction in Draper expression in ensheathing glia. The dotted box highlights the brain region displayed in C. M) Confocal images of the antennal lobe region of flies expressing either Luciferase-RNAi or Fis1-RNAi specifically in their adipose tissue immunostained with anti-Draper. N) The mean fluorescent intensity of Draper was measured within a region of interest (white box) that coincided with the location of the ensheathing glia. These measurements were derived from a Z-stack summation projection covering the entire depth of the antennal lobe. Student t-test with Welch’s correction. N = each circle represents an individual fly. O) Schematic for experimental design and summarized results of P and Q. Knocking down the antioxidant Sod2 in adipose tissue leads to an increase in Draper expression in ensheathing glia. The dotted box highlights the brain region displayed in C. P) Confocal images of the antennal lobe region of flies expressing either Luciferase-RNAi or Sod2-RNAi specifically in their adipose tissue immunostained with anti-Draper. Q) The mean fluorescent intensity of Draper was measured within a region of interest (white box) that coincided with the location of the ensheathing glia. These measurements were derived from a Z-stack summation projection covering the entire depth of the antennal lobe. Student t-test with Welch’s correction. N = each circle represents an individual fly. R) Schematic for experimental design and summarized results of S and T. Overexpression of the antioxidant Catalase in adipose tissue leads to a reduction in Draper expression in ensheathing glia. The dotted box highlights the brain region displayed in C. S) Confocal images of the antennal lobe region of flies overexpressing either Luciferase or Catalase specifically in their adipose tissue immunostained with anti-Draper. T) Mean fluorescent intensity of Draper measured within a region of interest (white box) that coincides with the location of ensheathing glia. These measurements were derived from a Z-stack summation projection covering the entire depth of the antennal lobe. Student t-test with Welch’s correction. N = each circle represents an individual fly.

Under starved conditions, glia utilizes ketone bodies as an alternative energy source for mitochondrial oxidative phosphorylation. Consequently, β-HB shifts brain metabolism from glycolysis to oxidative phosphorylation^86^. Since glial activation requires a metabolic shift from oxidative phosphorylation to glycolysis, we hypothesized that ketone bodies would deactivate glia, reducing Draper expression. This hypothesis aligns with our previous findings that a high-sugar diet (HSD) induces a metabolic shift in the brain towards oxidative phosphorylation and away from glycolysis^53^. To determine if adipocyte-produced ketone bodies impact glial Draper expression, we performed RNAi-mediated knockdown of the ketogenesis enzyme *Acat1* using an adipocyte-specific promoter. Disrupting ketone body production in adipocytes significantly increased glial Draper levels, suggesting that adipocyte-derived ketone bodies negatively regulate glial Draper expression (Figure 3H-J).

Mitochondrial morphology dictates function, with elongated mitochondria typically associated with reduced ROS. We specifically knocked down the mitochondrial fission protein-encoding gene *Fis1* in adipose tissue to investigate whether HSD-induced mitochondrial elongation affects glial Draper levels. Flies with *Fis1* knockdown showed reduced Draper levels in their ensheathing glia compared to those with the control (*Luciferase-RNAi*) (Figure 3L-N). We next assessed if adipocyte ROS levels impact Draper signaling. Given that HSD-fed flies exhibited reduced oxidized mitochondria and elevated antioxidants (Figure 2I-K), we reasoned that increasing adipocyte ROS would upregulate Draper levels. Indeed, knocking down the mitochondrial antioxidant Superoxide dismutase 2 (*Sod2*) in adipose tissue led to increased glial Draper expression (Figure 3O-Q). Conversely, overexpressing the antioxidant *Catalase* in adipose tissue reduced Draper levels in ensheathing glia, similar to HSD-fed flies (Figure 3R-T).

These findings suggest that adipocyte mitochondrial metabolism and redox state regulate glial Draper levels. Overall, our results suggest that adipocyte lipid and mitochondrial metabolism remotely influences the glial phagocytic state.

### Adipocyte-derived ApoB lipoprotein Signals regulate Glial Phagocytic Response to Neuronal Injury

One of the most well-known nutrient-sensitive adipokines is leptin and its *Drosophila* ortholog upd2. Released from adipose tissue in response to nutrient availability, upd2 functions as a satiety signal by binding to its JAK/STAT receptor, Domeless, in the brain^55,87^. Since JAK/STAT signaling in the ensheathing glia has been known to maintain Draper levels during axonal injury^88^, we investigated whether the adipokine upd2 – a JAK/STAT ligand^89^-remotely regulates the glial phagocytic state. We hypothesized that if this were the case, then loss of *upd2* would result in Draper downregulation like that observed with HSD and adipocyte-specific mitochondrial disruption. To test this, we measured Draper levels in the ensheathing glia of *upd2* deletion mutants. We found that loss of *upd2* did not affect Draper levels, suggesting that the adipo-glial coupling we observe is unlikely to be mediated by upd2/leptin (Figure S3) but by another adipocyte-derived signal that supports the glial phagocytic function.

As the body’s primary lipid repository, adipose tissue distributes lipids and fatty acids to various organs, including the brain. ApoB-Lpps (lipophorins) are primary carriers of adipocyte-derived lipids in circulation. Orthologous to mammalian apolipoprotein B, ApoB in flies is primarily expressed in adipose tissue and undergoes post-translational cleavage to produce two fragments, ApoI and ApoII, with ApoII containing the lipid-binding domain of the ApoB protein. We previously showed that prolonged HSD treatment reduces the amount of ApoB delivered to the brain^39^ (Refer to Schematic Figure 4A). Supporting this, we found that hemolymph concentrations and composition of PE phospholipids, the major component of the *Drosophila* lipoproteins—lipophorins^31^, are reduced in HSD-fed flies (Figure S4A-B). Given the dysregulation in adipocyte lipid homeostasis caused by HSD (Figure 2), we hypothesized that the HSD-induced disruption in glial Draper signaling could result from reduced ApoB signaling from adipocytes to glia. To test this, we performed RNAi-mediated knockdown of *ApoB* (*Apolpp*) using an adipocyte-specific promoter, then measured Draper levels in glia. *ApoB-RNAi* flies had reduced glial basal Draper levels (Figure 4C-E). Following neuronal injury, ensheathing glia typically upregulates Draper to enhance their ability to detect and engulf neuronal debris, preventing inflammation and secondary neuronal death. We asked whether ApoB regulates injury-induced Draper upregulation in ensheathing glia. To answer this, we performed unilateral antennal ablation and measured Draper levels one day later. Intriguingly, similar to HSD-fed flies ^53^, flies with adipocyte-specific knockdown of ApoB showed no upregulation of Draper after neuronal injury (Figure 4E), suggesting that ApoB signaling from adipocytes positively regulates glial phagocytic status.

**Figure 4.**
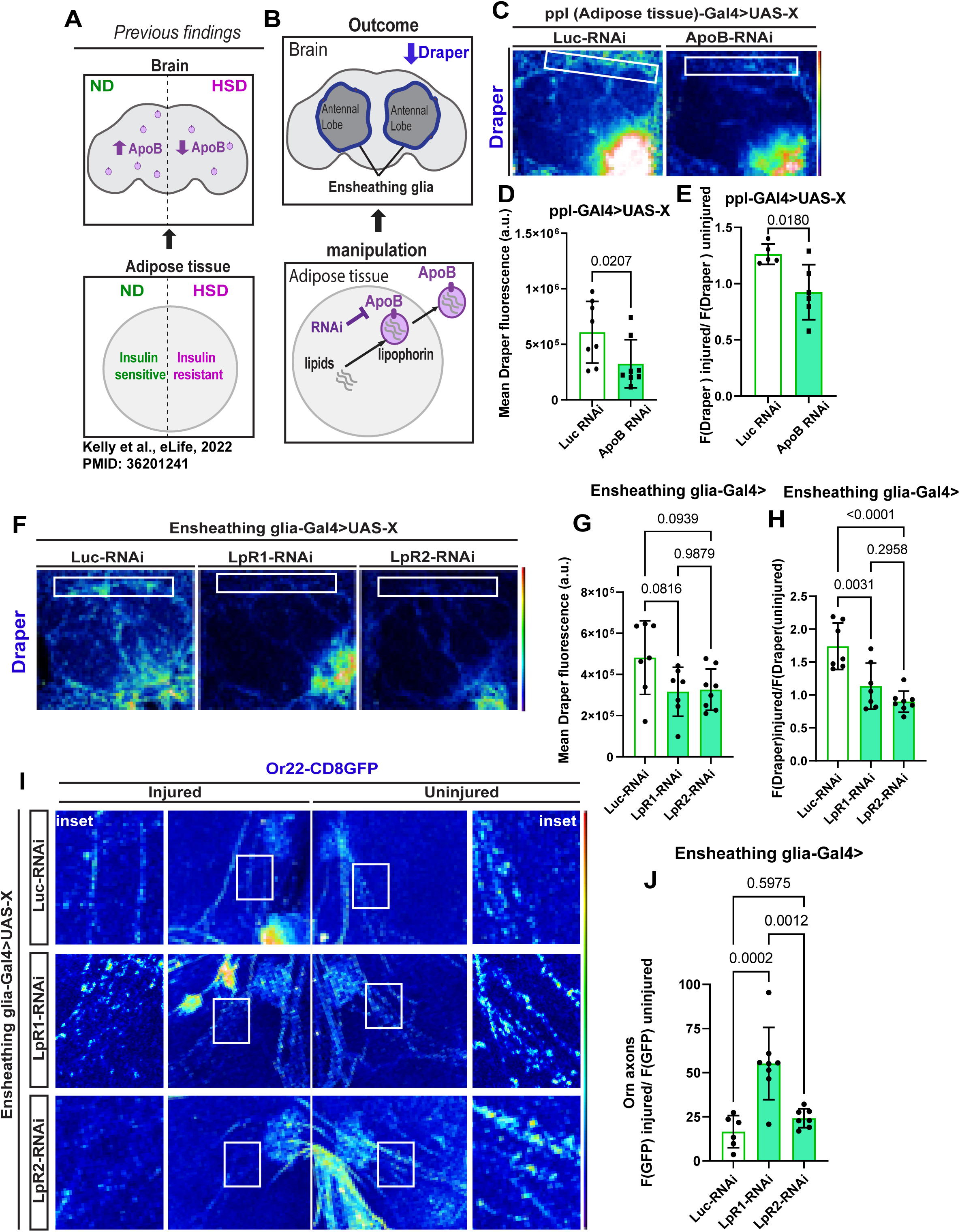
Disrupting adipocyte to glia apolipoprotein signaling diminishes Draper-mediated glial response to axonal injury. A) Schematic summarizing previously published results. Prolonged HSD exposure (2 weeks) leads to adipocyte insulin resistance, disrupting ApoB delivery to the brain. B) Schematic for experimental design and summarized results of C-E. Knocking down ApoB in adipose tissue leads to a reduction in Draper expression in ensheathing glia. The dotted box highlights the brain region displayed in C. C) Confocal images of the antennal lobe region of flies expressing either Luciferase-RNAi or ApoB-RNAi specifically in their adipose tissue immunostained with anti-Draper. D) Mean fluorescent intensity of basal Draper measured within a region of interest (white box) that coincides with the location of ensheathing glia. These measurements were derived from a Z-stack summation projection covering the entire depth of the antennal lobe. Student t-test with Welch’s correction. N = each circle represents an individual fly. E) Mean fluorescent intensity of Draper levels one-day post-injury, normalized to the uninjured control, was measured within a region of interest (white box) corresponding to the location of ensheathing glia. These measurements were obtained from a Z-stack summation projection covering the entire depth of the antennal lobe. Student t-test with Welch’s correction. N = each circle represents an individual fly. F) Confocal images of the antennal lobe region of flies expressing either Luciferase-RNAi, LpR1-RNAi, or LpR2-RNAi specifically in their ensheathing glia immunostained with anti-Draper. G) Mean fluorescent intensity of basal Draper measured within a region of interest (white box) that coincides with the location of ensheathing glia. These measurements were derived from a Z-stack summation projection covering the entire depth of the antennal lobe. Student t-test with Welch’s correction. N = each circle represents an individual fly. H) Mean fluorescent intensity of Draper levels one-day post-injury, normalized to the uninjured control, was measured within a region of interest (white box) corresponding to the location of ensheathing glia. These measurements were obtained from a Z-stack summation projection covering the entire depth of the antennal lobe. Student t-test with Welch’s correction. N = each circle represents an individual fly. I) 3D projections of the antennal lobe region of flies expressing membrane-tagged GFP in a subset of olfactory neurons (Orn22a) and expressing either Luciferase-RNAi, LpR1-RNAi, or LpR2-RNAi specifically in their ensheathing glia. J) Mean fluorescent intensity of GFP in injured axons normalized to uninjured axons within the same fly. Measurements were obtained from z-stack summation projections one day after unilateral antennal ablation. Analysis was conducted within a defined region of interest (white box) corresponding to the location of olfactory axons.

We previously showed that HSD causes ensheathing glial insulin resistance by stimulating systemic insulin release from insulin-producing cells (IPCs) in the brain, leading to impaired glial phagocytosis. Other adipocyte-derived lipophorins (Lipid Transfer Particle-LTP) regulate insulin-producing cells (IPCs)^38^. Hence, we investigated whether the adipocyte-specific ApoB knockdown affects IPCs by examining *Drosophila* insulin-like peptide 5 (Dilp5) levels, which serve as a readout for Dilp secretion. However, we found no significant effect on Dilp5 levels (Figure S5), indicating that the downregulation of ensheathing glial Draper is unlikely to be caused by ApoB-mediated insulin signaling dysregulation. These results suggest that adipocyte-to-brain ApoB-chaperoned lipid transfer regulates Draper-mediated glial response to neuronal injury.

The evolutionarily conserved low-density lipoprotein receptors LpR1 and LpR2^32^ facilitate the uptake of ApoB-chaperoned lipids. To determine whether glial Draper signaling downstream of ApoB is via ApoB-chaperoned lipophorins, we knocked down *LpR1* and *LpR2* using RNAi, specifically in the ensheathing glia. Upon measuring basal levels of Draper, we found that both LpR1 and LpR2 knockdowns resulted in reduced Draper levels, although the reduction was not statistically significant at baseline (Figure 4G-H). Strikingly, LpR1 and LpR2 knockdowns showed no upregulation of Draper after axonal injury (Figure 4I), like those with adipocyte-specific ApoB knockdowns (Figure 4F), suggesting that ApoB-chaperoned lipophorins help glial cells maintain phagocytic state post-injury.

To understand how adipocyte-glia ApoB signaling could impact glial phagocytic function, we knocked down LpR1 and LpR2 in the ensheathing glia of flies expressing membrane-tagged GFP in a subset of olfactory neurons (Odorant receptor 22a). We then performed unilateral antennal ablation and examined GFP accumulation one day later. We established an endogenous control by normalizing GFP fluorescence on the injured side to the uninjured side of the same animal. We found that LpR1 knockdown significantly impaired the ability of ensheathing glia to clear axonal debris, indicated by high levels of GFP accumulation. Surprisingly, LpR2 knockdown did not significantly impair glial phagocytic function compared to LpR1 knockdown (Figure 4J-K) despite causing similar disruptions in Draper signaling (Figure 4G-I). This suggests that pathways downstream of Draper are disrupted in LpR1 knockdown but are either unaffected or possibly compensated for in LpR2 knockdown (see discussion). These findings indicate that adipocyte-to-glia LpR1-mediated ApoB signaling supports glial phagocytic function by increasing Draper levels.

Altogether, we uncover a previously unappreciated role of adipose tissue in supporting glial phagocytic function via lipoprotein-mediated signaling. Importantly, we show that prolonged exposure to HSD disrupts this adipo-glial metabolic coupling, directly impacting glial phagocytic function. Mid-life obesity is an independent risk factor for cognitive decline ^1–4^, but the mechanisms by which adipocytes impact brain function are hard to pinpoint. Using a sophisticated genetic model system, we establish that the adipocyte metabolic state regulates glial biology via lipoprotein signals. Hence, our findings are directly relevant to managing and developing therapies for obesity-mediated cognitive decline.

## Discussion

The adipose tissue-brain axis is pivotal for brain function, with adipocytes serving as dynamic endocrine organs that secrete vital signaling molecules, including lipids and adipokines. In this study, we explored the intricate connection between adipocyte metabolism and glial function. Our findings reveal that high sugar diet (HSD)-induced obesity disrupts lipid homeostasis adipocytes, impairing glial function through a novel adipocyte-glial metabolic coupling. We identify Apolpp (ApoB) as an adipocyte-glia signal that regulates glial phagocytosis, a crucial process for maintaining neuronal health, via its receptor LpR1.

In the mammalian brain, a recent study by Fledrich and colleagues^90^, established that signaling between adipocytes and glial cells after nerve injury, with the adipokine leptin emerging as a key regulator of Schwann cell metabolism during the regenerative process. In our research, we provide compelling evidence that the metabolic status of *Drosophila* adipocytes regulates the baseline phagocytic activity of microglia-like cells—specifically, the ensheathing glia. However, we find that this adipocyte-glial metabolic communication in regulating the phagocytic state does not rely on *Drosophila’s* leptin homolog, Upd2, but is instead governed by ApoB-Lpps. Furthermore, this suggests that different glial cell types are likely regulated by a myriad of incoming signals. Taken together, these studies reveal an emerging paradigm in which adipocytes play a pivotal role in regulating and supporting glial function. Importantly, our study has shown that adipo-glial signals extend beyond traditional adipokine signaling and includes the involvement of ApoB-Lpp lipid carriers.

### Starvation amidst nutritional abundance: adipocyte metabolic adaptation in diet-induced insulin resistance

During starvation, adipocytes adapt by enhancing lipolysis, releasing fatty acids from lipid droplets to provide an alternative energy source, and increasing fatty acid oxidation (FAO) to compensate for reduced glycolysis. Interestingly, the metabolic adaptations observed in insulin resistance resemble the body’s response to starvation^91^. Consistent with this, we observed that prolonged exposure to HSD, decreased glycolytic enzymes like *pyk* and Ldh, reduced levels of the lipolysis inhibitor Plin1, and increased expression of key enzymes in fatty acid oxidation (FAO) in the adipose tissue of HSD-fed flies. While these adaptations support short-term energy balance under stress, sustained reliance on FAO can elevate circulating fatty acids and their metabolites^92,93^, potentially causing harmful lipotoxic effects in the brain. Indeed, we observed that prolonged HSD increases ketone body production, a hallmark of prolonged nutrient deprivation. Hence, prolonged HSD creates a starvation-like metabolic adaptation.

### Prolonged HSD alters the mitochondrial state in adult *Drosophila* adipocytes

Increased mitochondrial activity can produce excessive reactive oxygen species (ROS). Remarkably, fewer oxidized mitochondria are present after three weeks of HSD exposure, potentially due to increased antioxidant expression mediated by Nrf2 upregulation. This suggests prolonged HSD exposure may trigger a hormetic response, effectively reducing ROS levels, whereas shorter exposures may temporarily elevate ROS levels. Studies have shown obesogenic diet stress in mice can trigger compensatory antioxidant signaling in the heart tissue, which originates from adipocytes^94^. Another reason for HSD-induced antioxidant upregulation could be the expansion of lipid droplets in adipose tissue, which studies have shown act as protective sinks for ROS^95,96^.

Furthermore, upregulating fatty acid oxidation (FAO) enhances lipid droplet-mitochondria interaction to facilitate efficient fatty acid transfer to mitochondria. This heightened interaction reduces fatty acid oxidation in the cytosol, thereby mitigating oxidative stress^97,98^. Future work will be needed to determine whether reduced mitochondrial oxidative stress in adipocytes is communicated to the brain.

### The adipose tissue-brain axis controls glial response to neuronal injury

The primary carrier of lipids from the periphery to the brain is ApoB^19–22^. ApoB crosses the blood-brain barrier (BBB) by binding to its low-density lipoprotein receptor (LDLR) receptor, expressed by both glial cells and neurons^99^. Our previous study demonstrated that the delivery of *Drosophila* ApoB to the brain on HSD is reduced^39^. Notably, insulin signaling regulates the expression of LDLR, with insulin resistance leading to LDLR inactivation^100^. Since we have also shown that HSD causes glial insulin resistance, HSD likely affects ApoB signaling at both the adipocyte and the glial level. Knocking down the *Drosophila* LDLR, LpR, in neurons induces neurodegeneration^101^, highlighting an ApoB-LpR mediated mechanistic link between obesity and neurodegeneration.

### Distinct pathways regulate basal and injury-induced Draper expression

Phosphoinositide 3-kinase (PI3K) controls basal Draper expression, while the Janus kinase/signal transducer and activator of transcription (JAK/STAT) pathway governs injury-induced Draper expression^88^. Interestingly, LDL receptor (LDLR) signaling influences both pathways. Our findings support this, demonstrating that adipocyte ApoB knockdown reduces basal and injury-induced Draper upregulation. However, knockdown of the *Drosophila* homologs of LDLR, LpR1, and LpR2 does not significantly affect basal Draper levels but impairs injury-induced Draper upregulation. This suggests that LpRs may serve redundant functions under normal conditions but are essential for injury-induced Draper upregulation, attributable to increased energetic demands of glial cells during phagocytosis. Intriguingly, while LpR1 knockdown accumulates neuronal debris post-injury, indicating glial phagocytic impairment, knocking down LpR2 does not impair glial phagocytosis. LpR1 and LpR2 play overlapping but distinct roles in specific contexts, such as during the delivery of lipoproteins to the ovary^32^. This leads us to surmise that LpR1 is predominant in injury-induced glial response to neurite debris clearance. Future work is needed to delineate how signaling downstream of LpR1 leads to Draper expression.

### HSD-induced disruption in lipid homeostasis mimics the changes seen with aging

Several studies reveal changes in lipid homeostasis in the aging brain, specifically noting a marked accumulation of lipids in glial cells. This accumulation is beneficial as it sequesters cytotoxic peroxidated lipids from neurons ^37,102^. We have previously shown that HSD-fed fly brains exhibit increased lipid droplet accumulation^53^, similar to the aging brain. This may seem counterintuitive, given that HSD disrupts ApoB delivery to the brain, presumably carrying lipids from adipocytes. However, Palm et al. demonstrated that knocking down ApoB reduced medium-chain diacylglycerols (DAGs), not long-chain DAGs^31^. This distinction is intriguing because long-chain DAGs are synthesized endogenously in the brain, while medium-chain DAGs are sourced from the diet^103^. Therefore, the brain may upregulate endogenous lipid synthesis to compensate for HSD-induced disruption in ApoB-mediated delivery of lipids from adipocytes. Future studies will investigate how adipocyte-brain ApoB signaling regulates the glial lipidome and influences glial phagocytic function.

### ApoB-LpR1 Draper signaling and possible mechanisms via lipid-based signaling

Our studies show that ApoB-Lpp signaling as well as adipocyte metabolic state directly impacts Draper expression in ensheathing glia, that in turn maintains its phagocytic competence. Future work is required to tease out the mechanism by which ApoB-Lpp increase Draper expression. Nakanishi and Colleagues have shown in independent studies that Draper is a multi-liganded phagocytic receptor^104–106^. Studies from Freeman and colleagues have shown that Draper upregulates its own expression by an auto-regulatory positive feed forward loop^88^. Taken together these suggest an intriguing possibility is that ApoB-Lpps could ligand and activate Draper expression.

The extracellular region of Draper contains three conserved, cysteine-rich domains— EMI, NIM, and EGF-like—that are common to many proteins. The EMI and NIM domains are located near the N-terminus bind to phosphatidylserine (PS), an “eat-me” signal essential for processes like phagocytosis^104^; the authors also found that Draper binds to phosphatidylethanolamine (PE)-rich liposomes^104^, but not to phosphatidylcholine (PC)-rich species. ApoB transports PE-rich lipids to the brain^31^. Our recent work^39^ further revealed that disrupting PE production in fat tissue, particularly under a high-sugar diet (HSD), impairs the delivery of PE-rich ApoB-lipoprotein particles (ApoB-Lpp) to the brain. Hence, an attractive hypothesis is that ApoB-Lpp delivers PE-rich lipid ligands to Draper’s EMI and NIM domains, activating Draper signaling via auto-regulatory loop, and would require further testing.

Alternatively, but not mutually exclusive hypothesis, is that LpR1 may regulate Draper expression, in response to injury and at baseline. For instance, activity dependent upregulation of LpR1 has been shown to play a role in dendrite morphogenesis in *Drosophila* larvae^107^. In a similar vein, LpR1 may be liganded by ApoB at baseline, and upregulated in response to injury, to potentiate Draper signaling via increasing its transcription. Future work will be needed to test these hypotheses and to understand how Draper expression is regulated by ApoB-LpR axis.

In conclusion, this study uncovers that the lipid homeostasis and mitochondrial metabolism of adipocytes directly influence the phagocytic activity of glial cells through ApoB lipoprotein-mediated communication. Independent human studies have consistently demonstrated a strong correlation between mid-life obesity and cognitive decline in later life. Despite this, pinpointing the causal links between adipocyte metabolic states and brain health remains challenging. By revealing that adipocyte metabolic status directly impacts glial function in *Drosophila*, our findings offer critical insights that inform how to manage and treat brain disorders stemming from obesogenic conditions.

## Methods

### Fly husbandry

The following *Drosophila* strains were used: *Or22a-mCD8GFP* (BDSC #52620), *Ensheathing glia-Gal4* (BDSC # 39157), *UAS-Plin1::GFP*^108^, Ldh-GFP (a generous gift from Dr. Tennessen), *ppl-Gal4*^55^, *Lpp-Gal4*^109^, *UAS-mitoTIMER* (BDSC#57323), UAS-Fis1-RNAi (BDSC# 63027), *UAS-SOD2-RNAi* (BDSC#36871), *UAS-Apolpp-RNAi* (BDSC# 33388), *UAS-LpR1-RNAi* (BDSC# 27249), *UAS-LpR2-RNAi* (BDSC# 31150), UAS-Catalase (BDSC# 24621), *UAS-whd-RNAi* (BDSC# 34066), *UAS-Acat1-RNAi* (BDSC# 51785), *UAS-Luc-RNAi* (BDSC#31603). Flies were housed in 25°C incubators, and all experiments were done on 7-10-day-old adult male flies. The flies’ diet consisted of 15 g yeast, 8.6 g soy flour, 63 g corn flour, 5 g agar, 5 g malt, and 74 mL corn syrup per liter.

### Antennal nerve injury

As adapted from^8,47–50,53,88^, flies were anesthetized using CO2, and antennal nerve injury was accomplished by unilaterally removing the third antennal segment of anesthetized adult flies using forceps. Flies were then placed back into the vial until they were dissected 24 hours after injury.

### Immunostaining

Immunostaining of adult brains and fat bodies was performed as previously described [65,66]. Tissues were dissected in ice-cold PBS. Brains were fixed overnight in 0.8% paraformaldehyde (PFA) in PBS at 4°C. The fixed brains were washed 5 times in PBS with 0.5% BSA and 0.5% Triton X-100 (PAT), blocked for 1 h in PAT + 5% NDS, and then incubated overnight at 4°C with the primary antibodies. Following incubation, the brains were washed 5 times in PAT, re-blocked for 30 min, and then incubated in secondary antibody in block for 4 h at room temperature. Finally, the brains were washed 5 times in PAT, then mounted on slides in Slow fade gold antifade. Primary antibodies were as follows: Chicken anti-GFP (1:500; Cat# ab13970, RRID:AB_300798), Mouse anti-Draper (1:50; DSHB 5D14 RRID:AB_2618105), Rabbit mitomCherry (1:500; Cat#ab167453, RRID:AB_2571870). Secondary antibodies from Jackson ImmunoResearch (1:500) include donkey anti-Chicken Alexa 488 (Cat# 703-545-155, RRID: AB_2340375), donkey anti-rabbit Alexa 594 (Cat# 711-585-152, RRID: AB_2340621), and donkey anti-mouse Alexa 594 (Cat# 715-585-150, RRID: AB_2340854).

### Image analysis

Images were acquired with a Zeiss LSM 800 and Leica Stellaris confocal system. Images were analyzed using ImageJ. All images within each experiment were acquired with the same confocal settings. Z-stack summation projections at 0.5 μm intervals were generated, and a region of interest (indicated in the fig) was used to measure the integrated density values of each fluorescent tag. A maximum-intensity projection of Z-stacks that covered the full depth of the antennal lobe was used for ImageJ analysis to measure mitochondrial morphology. The maximum-intensity projection was inverted and automatically thresholded before the ‘analyze particles’ function was used to measure the average mitochondrial circularity major and minor axes. The size and dimensions of all ROIs were maintained consistently throughout each experiment. Dilp 5 levels were quantified using z-stack summation projections to capture the full depth of the IPCs. A region of interest (ROI) around the IPCs was manually outlined with the freehand tool, and integrated density values were then measured.

### Hemolymph extraction

The protocol for adult fly hemolymph extraction was adapted from. Briefly, 30 adult flies were anesthetized on a CO2 pad and punctured in the thorax region with a tungsten needle. The flies were then transferred into a 0.5 mL Eppendorf tube with 5 holes made in the bottom using an 18G needle. This 0.5 mL tube was placed inside a 1.5 mL Eppendorf tube containing 30 µL of PBS and centrifuged at 5000 RPM for 5 minutes. The samples were then flash-frozen in liquid nitrogen until ready to use.

### Sample Prep for metabolomics analysis

Aqueous metabolites for targeted LC-MS profiling of whole flies and hemolymph samples were extracted using a protein precipitation method similar to the one described elsewhere.

#### Whole fly Samples

Whole adult male flies were frozen in liquid nitrogen after 7 or 14 days on a normal diet or HSD. Ten flies were used per biological sample, and 3 biological replicates were used for each diet and time point. Samples were first homogenized in 200 µL purified deionized water at 4 °C. Then 800 µL of cold methanol containing 124 µM 6C13-glucose and 25.9 µM 2C13-glutamate was added (reference internal standards were added to the samples to monitor sample prep). Afterward, samples were vortexed, stored for 30 minutes at −20 °C, sonicated in an ice bath for 10 minutes, centrifuged for 15 min at 14,000 rpm and 4 °C, and then 600 µL of supernatant was collected from each sample. Lastly, recovered supernatants were dried on a SpeedVac and reconstituted in 0.5 mL of LC-matching solvent containing 17.8 µM 2C13-tyrosine and 39.2 3C13-lactate (reference internal standards were added to the reconstituting solvent to monitor LC-MS performance). Samples were transferred into LC vials and placed into a temperature-controlled autosampler for LC-MS analysis.

#### Hemolymph Samples

30 flies were used per biological sample, and 3 biological replicates were used for each diet and timepoint. Samples were thawed at 4 °C for 60 mins and vortexed for 10 sec. 50uL of each sample was transferred to a 2 mL Eppendorf tube, and 50 uL of 50%MeOH/50%Water containing 30 stable isotope-labeled internal standards (SILISs) was added. Afterward, 250 uL of cold MeOH containing two additional SILISs was added to each sample. Samples were vortexed, stored for 30 minutes at −20 °C, centrifuged for 15 min at 14,000 rpm and 4 °C, and then 250 µL of supernatant was collected from each sample. Lastly, recovered supernatants were dried on a SpeedVac and reconstituted in 0.5 mL of LC-matching solvent containing two more SILISs. 34 SILISs were added to the samples in various sample prep steps to monitor sample prep, assay performance, and determine absolute concentrations for the metabolites that had corresponding SILISs.

### Metabolomics

Targeted LC-MS metabolite analysis was performed on a duplex-LC-MS system composed of two Shimadzu UPLC pumps, CTC Analytics PAL HTC-xt temperature-controlled auto-sampler, and AB Sciex 6500+ Triple Quadrupole MS equipped with ESI ionization source (2). UPLC pumps were connected to the auto-sampler in parallel and could perform two chromatography separations independently from each other. Each sample was injected twice on two identical analytical columns (Waters XBridge BEH Amide XP), where separations were performed in hydrophilic interaction liquid chromatography (HILIC) mode. While one column was performing separation and MS data acquisition in ESI+ ionization mode, the other column was getting equilibrated for sample injection, chromatography separation, and MS data acquisition in ESI-mode. Each chromatography separation was 18 minutes (total analysis time per sample was 36 minutes). MS data acquisition was performed in multiple-reaction-monitoring (MRM) mode. LC-MS system was controlled using AB Sciex Analyst 1.6.3 software. Measured MS peaks were integrated using AB Sciex MultiQuant 3.0.3 software. Up to 158 metabolites (plus 4 spiked standards) were measured across the fly samples study set, and up to 148 metabolites (plus 34 SILISs) were measured in the hemolymph sample set. For the hemolymph set, absolute concentrations of 30 metabolites were determined. In addition to the two study samples, two sets of quality control (QC) samples were used to monitor the assay performance and data reproducibility. One QC [QC(I)] was a pooled human serum sample used to monitor system performance, and the other QC [QC(S)] was pooled study samples. This QC was used to monitor data reproducibility. Each QC sample was injected per every 10 study samples. The median CV for the fly set was 2.8%, while for the blood samples was 11.1%.

### Lipidomics

Frozen hemolymph samples from ND and HSD-fed flies were sent to the Northwest Metabolomics Research Center for targeted quantitative lipid profiling using the Sciex 5500 Lipidyzer. Detailed methods can be found here^110^. The materials used include LC-MS grade methanol, dichloromethane, and ammonium acetate, all sourced from Fisher Scientific (Pittsburgh, PA). HPLC grade 1-propanol was obtained from Sigma-Aldrich (Saint Louis, MO). Milli-Q water was produced using an in-house Ultrapure Water System by EMD Millipore (Billerica, MA). The Lipidyzer isotope-labeled internal standards mixture, which contained 54 isotopes from 13 different lipid classes, was acquired from Sciex (Framingham, MA).

### qPCR

Twenty were dissected in RNAlater, then placed in 30 μl of TriReagent and a scoop of beads in a 1.5 mL safelock tube. The heads were homogenized using a bullet blender. RNA was then isolated using a Direct-zol RNA microprep kit following the manufacturer’s instructions. Isolated RNA was synthesized into cDNA using the Bio-Rad iScript RT supermix for RT-qPCR, and qPCR was performed using the Bio-Rad ssoAdvanced SYBR green master mix. Primers used are as follows: alpha-tubulin (endogenous control), forward: AGCGGTAGTGTCTGCCGTGT and reverse: CCAGCGTGGATTTGACCGGA; hnf4, forward: GGCGACGGGCAAACATTATG and reverse: CGCAAATCTGCAAGTGTACTGAT; cpt1, forward: TTGCCATCACCCATGAGGG and reverse: CAATCGCTTTTTCCAGGAACG; acc, forward: TGGCTGGTAGAGGAGAAGGG and reverse: CGTCTCACGGAGTTCACGTT; acat1, forward: ATTGCGAAGACCGATGTCCAG and reverse: GCAGCATACATTGGTGGGC; mcad, forward: TGGCACCTCTTTCGCACTC and reverse: GATGATCTCCTCACGGGTGAA; yip2, forward: TCTGCCGCAACCAAAGGTATC and reverse: GCGATCACATTTCCCACGATG; fis1, forward: GTCTGGCTTAAAATACTGCCGA and reverse: CATACCCTTTGCCACTTCCTT; opa1 forward: CGAGGAGTTCCTACTTGCCG and reverse: GTATCGCAGCTTGAGGGCTC; nrf2, forward: GAATGACCGCCGATCTCTTGG and reverse: GGAGCCCATCGAACTGACA. Relative mRNA quantification was performed using the comparative CT method and normalized to alpha-tubulin mRNA expression. Three technical replicates were used for each gene.

### Statistics

All statistical analyses were conducted using GraphPad PRISM (GraphPad Software Incorporated). Data are expressed as the mean ± standard deviation (SD). Specific statistical tests and sample sizes for each experiment are detailed in the figure legends. The normality of the data was assessed using the Shapiro-Wilk test. A two-tailed unpaired t-test with Welch’s correction or a two-way ANOVA with Holm-Sidak correction was applied for parametric analyses. For non-parametric analyses, a two-tailed t-test with Mann-Whitney correction was used. Statistical significance was defined as p < 0.05.

## Acknowledgments

We thank Dr. Jason M Tennessen for generously donating the *Ldh-GFP* transgenic fly line used in this article. We also thank Dr. Ava Brent and Zach Goldberg for their help with preparing the lipidomic and metabolomic samples. We thank Rajan Lab members –Dr. Kevin P. Kelly and Dr. Aditi Madan— for their critical reading, discussions, and comments on the manuscript. We acknowledge The Northwest Metabolomics Research Center at the University of Washington, Seattle, and NIH grant # 1S10OD021562-01 that purchased an LC-MS system for collecting targeted metabolic profiling data. Stocks obtained from the Bloomington Drosophila Stock Center (NIH P40OD018537) were used in this study.

---

**Figure S1.**
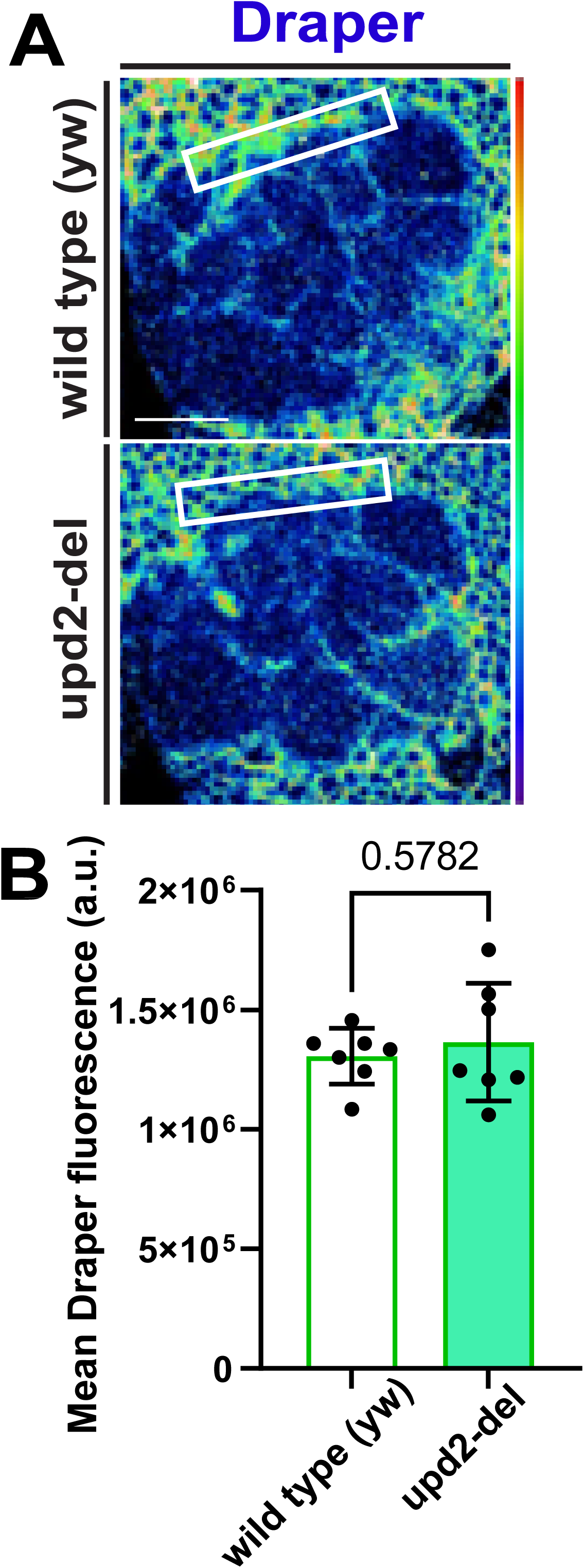
Adipocyte glycolytic breakdown occurs before it does in the brain. A) Confocal images of Ldh-GFP in the adipose tissue of flies fed either an ND or an HSD for 2 weeks. B) Mean Ldh-GFP fluorescent intensity, obtained from Z-stack summation projections of adipose tissue in ND and HSD-fed flies. Student’s t-test with Welch’s correction. N = each circle represents an individual fly. C) Timeline of Ldh reduction in adipose tissue versus the brain on HSD. Ldh is downregulated in adipose tissue prior to its downregulation in the brain.

**Figure S2.**
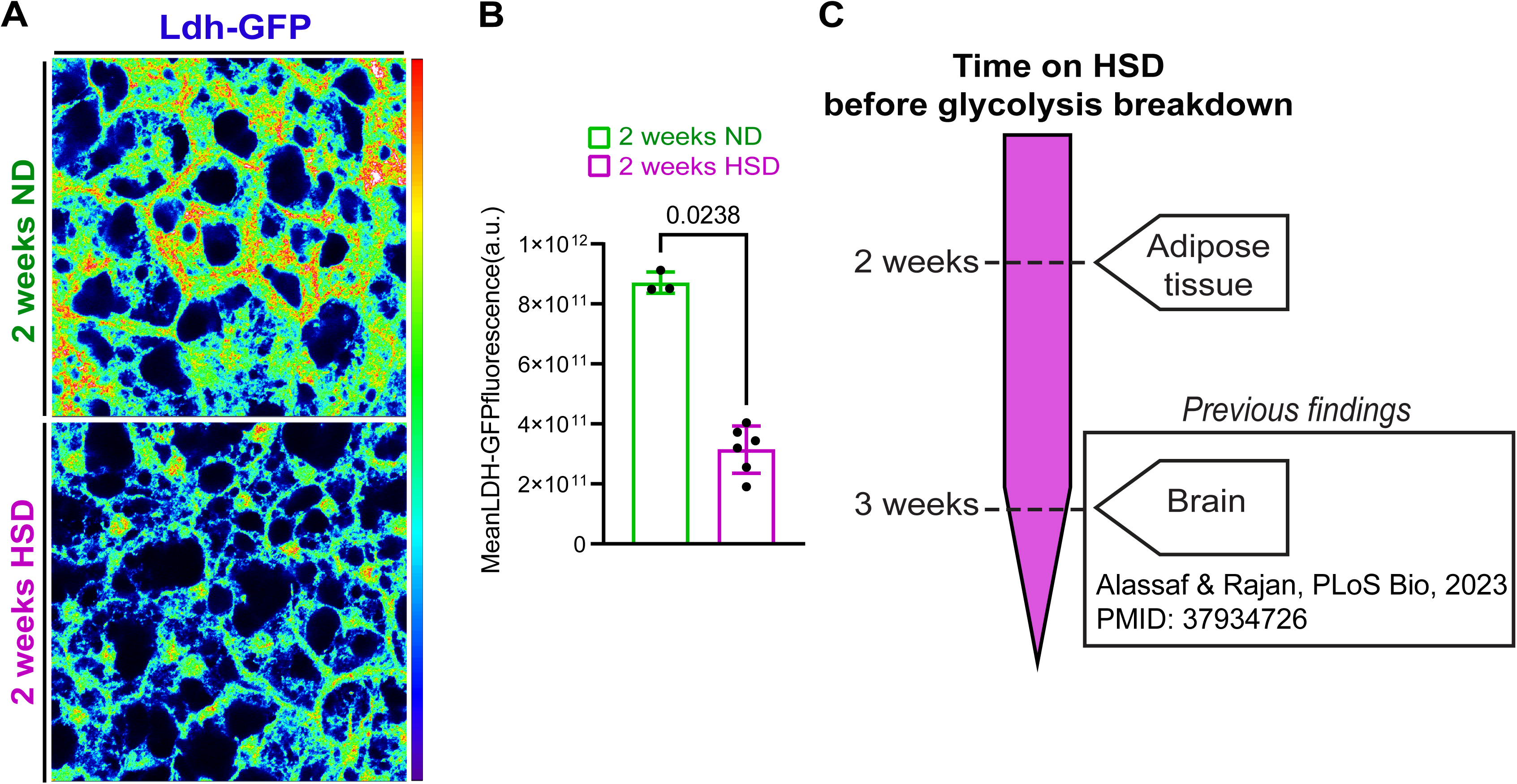
Adipocyte glycolytic breakdown occurs before it does in the brain. Mean beta-hydroxybutyrate in the hemolymph flies fed either an ND or an HSD for 1 week. Student’s t-test with Welch’s correction. N = 3 technical replicates of 30 flies/replicate.

**Figure S3.**
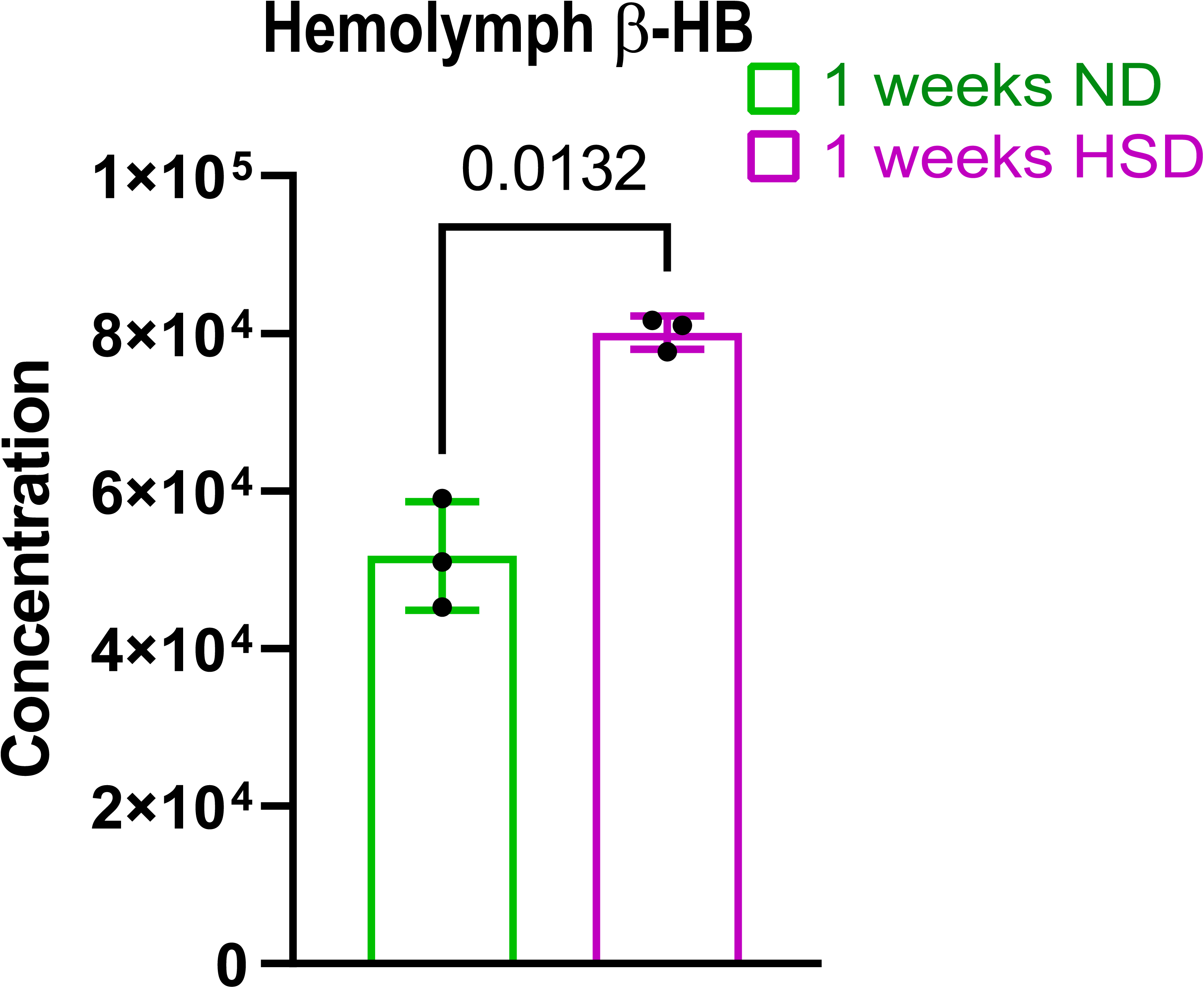
Upd2 does not affect Draper levels in the ensheathing glia. A) Confocal images of the antennal lobe region of wild type and upd2-deletion flies immunostained with anti-Draper. B) The mean fluorescent intensity of Draper was measured within a region of interest (white box) that coincided with the location of the ensheathing glia. These measurements were derived from a Z-stack summation projection covering the entire depth of the antennal lobe. Student t-test with Welch’s correction. N = each circle represents an individual fly.

**Figure S4.**
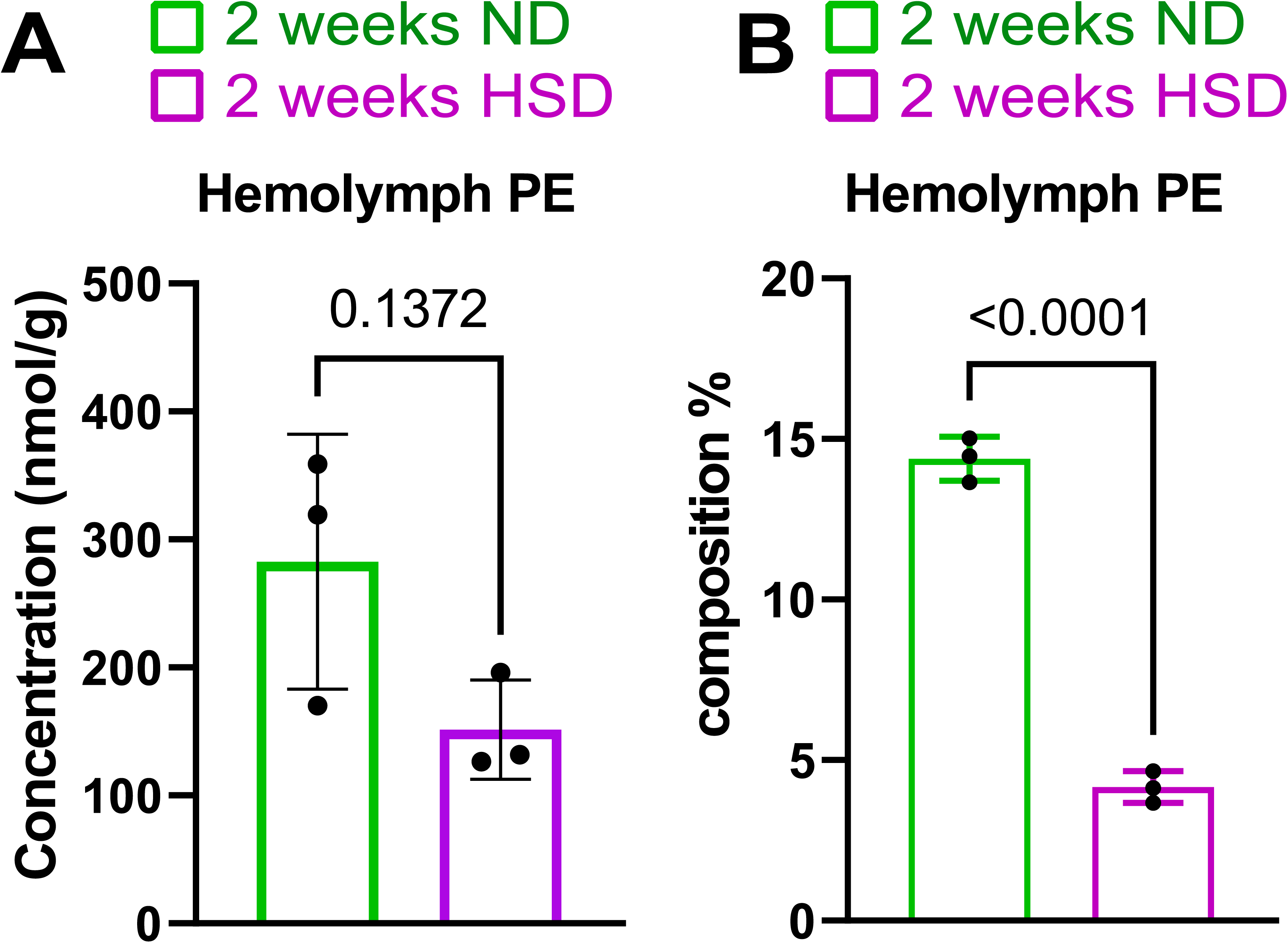
HSD treatment reduces hemolymph PE levels. A) The mean concentration of phosphatidylethanolamine (PE) in the hemolymph of flies fed either an ND or a HSD for 2 weeks. N = 3 technical replicates of 30 flies/replicate. B) The mean composition of phosphatidylethanolamine (PE) in the hemolymph of flies fed either an ND or a HSD for 2 weeks. N = 3 technical replicates of 30 flies/replicate.

**Figure S5.**
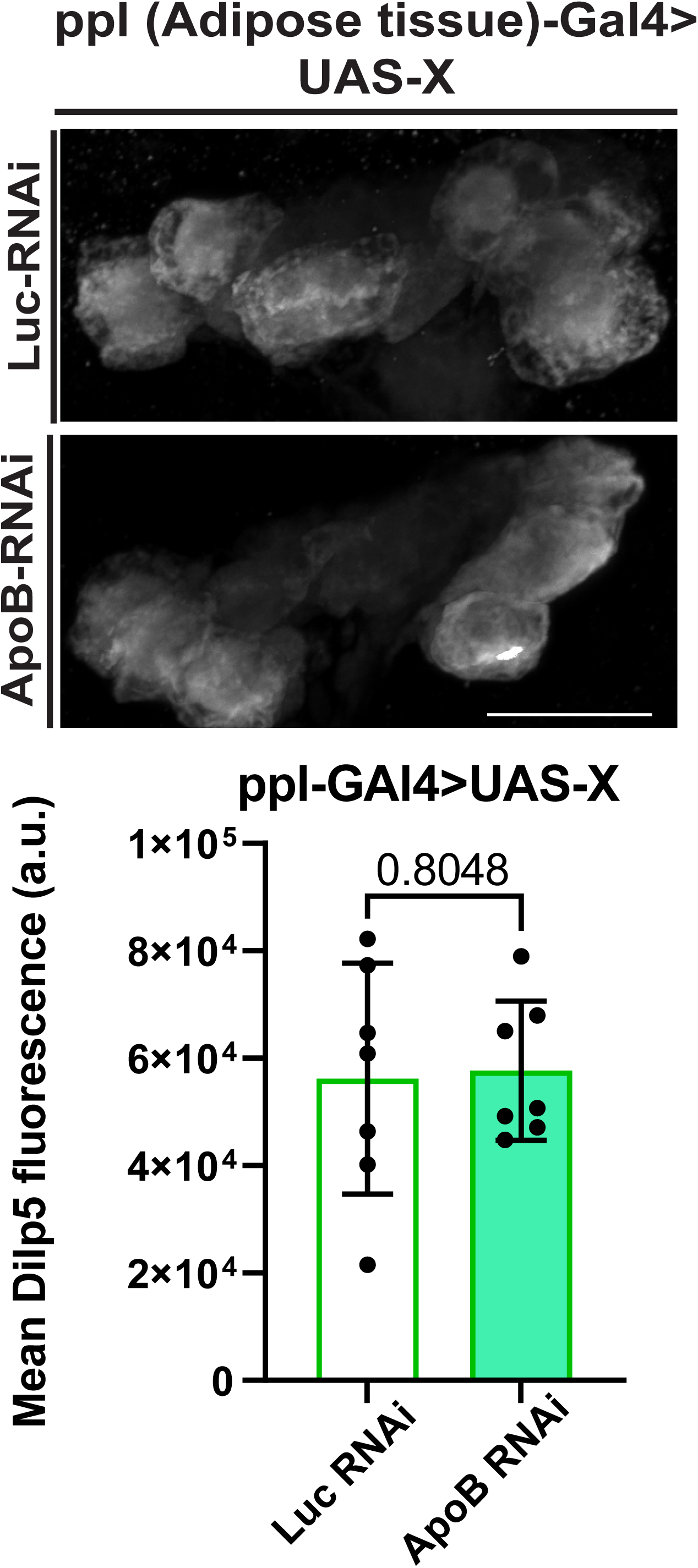
Knocking down ApoB in adipose tissue does not impact insulin levels in the brain. A) Z-stack summation projections of the insulin-producing cells (IPC) immunostained with anti-Dilp5 of flies expressing either Luciferase-RNAi or ApoB-RNAi specifically in their adipose tissue. B) Mean fluorescent intensity of Dilp5 derived from a Z-stack summation projection covering the entire depth of the IPCs. Student t-test with Welch’s correction. N = each circle represents an individual fly.

